# Proprioception is subject-specific and improved without performance feedback

**DOI:** 10.1101/850727

**Authors:** Tianhe Wang, Ziyan Zhu, Inoue Kana, Yuanzheng Yu, Hao He, Kunlin Wei

## Abstract

Accumulating evidence indicates that the human’s proprioception map appears subject-specific. However, whether the idiosyncratic pattern persists across time with good within-subject consistency has not been quantitatively examined. Here we measured the proprioception by a hand visual-matching task in multiple sessions over two days. We found that people improved their proprioception when tested repetitively without performance feedback. Importantly, despite the reduction of average error, the spatial pattern of proprioception errors remained idiosyncratic. Based on individuals’ proprioceptive performance, a standard convolutional neural network classifier could identify people with good accuracy. We also found that subjects’ baseline proprioceptive performance could not predict their motor performance in a visual trajectory-matching task even though both tasks require accurate mapping of hand position to visual targets in the same workspace. Using a separate experiment, we not only replicated these findings but also ruled out the possibility that performance feedback during a few familiarization trials caused the observed improvement in proprioception. We conclude that the conventional proprioception test itself, even without feedback, can improve proprioception but leave the idiosyncrasy of proprioception unchanged.

## Introduction

Knowing the spatial position of one’s hand is important for humans to maintain postures and perform actions. Both visual and proprioceptive cues are used for locating hands in space (Welch 1986; Van Beers et al. 1999). Though visual information plays a dominant role when both types of cues are available (Jeannerod 1988, 1991; Helms Tillery et al. 1994), proprioception continuously updates the nervous system about the hand location. It has been found that the hand location, if informed by proprioception alone, gradually drifts without visual calibration (Wann and Ibrahim 1992; Brown et al. 2003b, a). However, how proprioception changes over time has not been systematically investigated.

Previous studies have revealed that the accuracy of proprioception varies in the hand space, leading to spatial patterns of proprioceptive errors that are heterogeneous among individuals (van Beers et al. 1998; Haggard et al. 2000; Fuentes and Bastian 2009). On the group level, the accuracy of proprioception was affected by the distance from the body, with small proprioceptive errors in the areas close to the body and large errors in the areas away from the body (Wilson et al. 2010). The proprioceptive estimation of the left hand was biased to the left while that of the right hand to the right (Jones et al. 2010; Rincon-Gonzalez et al. 2011). Besides these general patterns on the group level, proprioception showed large inter-individual differences in the spatial pattern of accuracy (Brown et al. 2003b; Smeets et al. 2006). Measured by visual-matching tasks, the proprioception maps remained similar across conditions within a participant but differed widely across participants (Helms Tillery et al. 1994; Rincon-Gonzalez et al. 2011). As another indirect evidence of within-subject consistency, people also found that the proprioception map measured by a visual-matching task and by a pointing task were strongly correlated within a participant (Vindras et al. 1998).

However, to our knowledge, the subject-specificity of the proprioception map has never been systematically examined. Many previous studies reached their conclusions by eyeballing of data (Brown et al. 2003b; van den Dobbelsteen et al. 2004; Smeets et al. 2006). Other studies calculated the within-subject correlation coefficients between measurements from different conditions and found they were significantly larger than zero (Wann and Ibrahim 1992; Desmurget et al. 2000). However, this kind of correlation results only shows the similarity between conditions as opposed to the idiosyncrasy of proprioception maps between subjects. A couple of studies computed the within-subject correlation of proprioception maps and the between-subject correlation, but they did not compare these correlations, possibly due to a limited number of participants (Helms Tillery et al. 1994; Vindras et al. 1998; Rincon-Gonzalez et al. 2011). In sum, no previous study has quantitively examined the idiosyncrasy of the proprioception map, leaving the question open about to what extent one’s proprioception map can be distinguished from others’.

Proprioception underlies motor performance in various tasks (Rosenbaum 2009). Recent studies also found that motor learning and proprioceptive training could benefit each other if these two tasks were similar. Proprioceptive training by passively moving one’s hand around a target circle could improve the subsequent motor learning of drawing the target (Wong et al. 2012). Moreover, after a brief period of motor learning, i.e., tracing a series of visual targets, participants improved their accuracy of proprioception for more than 24 hours (Wong et al. 2011). Furthermore, the proprioceptive improvement was limited in the region where participants performed motor learning. With these findings, it is tempting to conjecture that proprioceptive capacity might be able to predict the motor performance of the tasks that require proprioceptive control of movements. A straightforward way to test this hypothesis is to examine the relationship between the baseline accuracy of proprioception and the baseline motor performance in the same workspace.

Here we used a hand visual-matching task with 100 target positions to obtain the proprioceptive error map in the reachable space. To quantitively study the subject-specificity of proprioception map across time, we measured proprioception multiple times over two days. To examine whether the baseline proprioceptive performance can predict motor performance, we then tested a trajectory production task that required accurate hand matching of visual templates. We found that the within-subject variance of proprioception errors was much smaller than the between-subject variance. Furthermore, based on people’s proprioception tested on the first day, a simple convolutional neural network classifier was able to identify the participant based on the proprioception map measured on the second day with an accuracy around 70% (base rate 1/47). We also found that proprioception measured by the visual-matching task could not predict the motor performance in the trajectory production task. Surprisingly, the accuracy of proprioception improved across days, even though our measurements did not provide performance feedback. In a separate experiment, we replicated our major findings and ruled out the possibility that limited performance feedback during the familiarization trials caused the improvement in proprioception across sessions.

## Methods

### Participants

A total of forty-seven graduate students and undergraduate students (30 males, age: 21.0 ± 2.2 yr, mean ± SD) of Peking University were recruited for two experiments, twenty-six for Experiment 1 and twenty-one for the Experiment 2. All participants were confirmed to be right-handed by the Edinburgh handedness inventory (Oldfield, 1971). All participants were new to the experimental task, naive to the purpose of the study, provided written informed consent before participating, and they received either course credit or monetary compensation for their time. All experimental protocols were approved by the Institutional Review Board of Peking University.

### Experimental setup

The experimental setup had been used in our previous researches (Yin and Wei 2014; Wei et al. 2014; Yin et al. 2016; Jiang et al. 2018). In all experiments, participants sat in front of a digitizing tablet and held the digital stylus with their left hand (Fig. 1A). They were instructed to match the tip of the stylus with either a point target or a trajectory target that was displayed on a horizontal display. The display was first projected on a back-projection screen horizontally placed above the tablet (LCD projector; Acer P1270, refreshing rate of 75Hz). The display was then reflected by a semi-silvered mirror placed horizontally at the chest level; the reflection matched in height with the tablet where the participant’s hand was. The participants viewed the stimulus and feedback in the mirror while their view of the hand and arm was occluded. The stylus movement on the tablet was one-to-one mapped onto the visual display after calibration. Participants were required to perform the location matching as accurate as possible with their preferred pace. They also centered their body with the tablet during the whole experiment. The task was controlled by a customized program written in MATLAB (Mathworks, Natick, MA; Psychophysics Toolbox).

**Fig.1.**
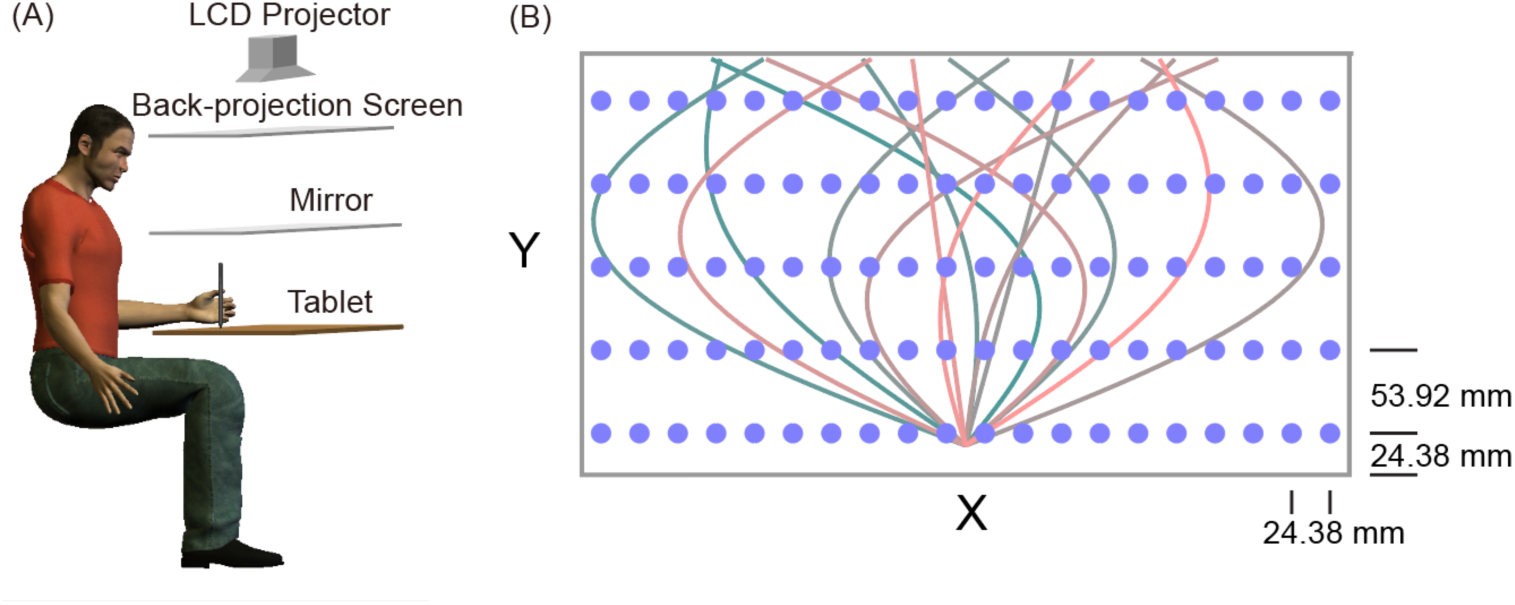
Experimental setup and material. A) Experimental setup. B) A schematic illustration of screen display during the experiment. Blue dots indicate the 100 target positions in the visual-matching task. Colored lines indicate the 15 target trajectories in the trajectory matching task.

### Tasks

#### Visual matching task

In each trial, a white light dot (50 mm diameter) was presented on the semi-silvered mirror to indicate the target position. The participants matched the target with the digital stylus held by the left hand. To obtain an accurate proprioception map, we included 100 target positions, which formed a 5 (row) × 20 (column) matrix in the workspace in front of the seated subject (Fig. 1B). The workspace was 48.76 cm wide and 26.96 cm long, located 20 cm in front of the seated participant. The distance between the adjacent columns was 24.38 mm, and that between the adjacent rows was 53.92 mm. Each target was measured once, and the order of targets was randomized. After the participants pressed the space bar of a keyboard with their right hand, the computer speaker played a beep sound to confirm the measurement. No performance feedback was given. The target disappeared directly while the next target appeared in a new position to start the next trial. The participants were allowed to move freely from one target position to the next at their own pace. Before formal data collection, we gave participants 16 familiarization trials for the visual matching task. Each trial was associated with a different target, and the 16 targets were evenly spaced to form a 4×4 matrix to cover the whole workspace. None of them overlapped with the targets in the formal test. For these familiarization trials, the actual position of the stylus was indicated by a green dot (50 mm diameter) for one second after the participant pressed the confirmation key. The 16 target positions were shown one by one from the bottom to the top and from left to right.

#### Trajectory matching task

The trajectory matching task was modified from a similar task in one of our previous studies (Dam et al. 2013). In the workspace of the visual matching task described above, we asked participants to produce a curved trajectory to “copy” a target trajectory that was visually presented on the projection screen (Fig. 1B). Each trial began with participants holding their left hand at a starting position indicated by a dashed circle (40 mm diameter) at the bottom center of the workspace. After 100 ms, the starting position changed from blue to green, and a beep sound was played to signal the incoming movement. Then, a target trajectory (a 20 mm-wide red line) appeared, stretching from the start position to the upper edge of the workspace. The target trajectory was prescribed by the formula: x = α × y + β × sin(πy), where y indicated the displacement in the depth direction and x indicated the displacement in the mediolateral direction. Numerically, y ranged between 0 and 1, where 1 represents 211 mm on the screen. Thus, the main direction and curvature of the curved trajectory were determined by α and β, respectively. Participants were instructed to make a fast movement to match the target trajectory without corrections accurately. During the movement, no cursor feedback was given to show their actual hand position. After reaching the upper edge of the workspace, another sound was played to indicate the end of the trial. The participant returned the stylus to the starting position without continuous cursor guidance. The hand location was only displayed as a white cursor (30 mm diameter) when it was within 5 cm around the starting position. No performance feedback was given, and a new trajectory appeared after one second.

To assess people’s performance for trajectory matching, we used fifteen target trajectories that were evenly distributed over the whole workspace (Fig. 1B). These trajectories were set by varying α from -1 to 1 and β from -0.9 to 0.8. All target trajectories started from the starting position at (x = 0, y = 0) and ended when y = 1. The target trajectories were presented in a random order, and each appeared twice in a row. Before the formal test, we gave each participant four trials to familiarize the task with a single target trajectory (α = 0, β = 0.1), which was not used in the formal experiment. In the first two practice trials, people received terminal feedback by viewing the actual movement trajectory made along with the target trajectory immediately after the movement end. The next two practice trials were the same as the formal trial without terminal feedback.

The participant was not allowed to start a movement before the start position turned green. Also, no backward movement towards the body was allowed. Warning messages, i.e., “Do not move before the start position turns green” or “Do not move backward,” were shown on the screen if these trials were detected. To avoid slow movement, we computed their average movement speed on each trial and compared it to the lowest speed allowed (165 mm/s). Movements slower than this threshold were regarded as invalid, and a warning message (“Too slow”) was displayed at the trial end to urge participants to move faster. All invalid trials were repeated immediately.

### Experimental protocols

#### Experiment 1

Experiment 1 was designed to test whether the proprioceptive performance was subject-specific and stable across days and whether it correlates to motor performance. It included three sessions with the first two sessions on day 1 and the third one on day 2 (Fig. 1C). There was a forty-minute rest between the first two sessions and a twenty-four hours interval between the last two sessions. The trajectory matching task was performed at the end of the first proprioception measurement, and it took about five minutes. Session 1 and session 3 started with a sixteen-trial familiarization, which provided participants with feedback at the end of each trial.

#### Experiment 2

We found that proprioception improved across sessions without any performance feedback in Experiment 1. One confound was that the 16 familiarization trials before session 3 provided performance feedback, which might improve people’s proprioceptive performance, as shown in the subsequent measurement sessions. In Experiments 2, we removed the familiarization trials before session 3 and added a 4th session with its own familiarization trials. Other procedures remained the same as in Experiment 1. Therefore, Experiment 2 included four sessions, two on the first day and two on the second day. We were particularly interested in the proprioceptive test of session 3: the previously observed improvement in this session should be absent if it was a result of familiarization trials with feedback. Similarly, we should observe an improvement in session 4 if the familiarization trials mattered.

### Data Analysis

The overall proprioception accuracy was quantified by the average visual-matching error at the 100 target positions in one session. The error was defined as the Euclid distance between the location of the target position (x_t_, y_t_) and the actual position of the stylus tip. To compare the proprioception error in different areas, we divided the workspace evenly into left and right regions by the vertical midline, and into inside and outside regions by the horizontal midline. Thus, the left region covered the ten columns of targets on the left, and the right region covered the other ten columns of targets on the right. The inside region covered the three rows close to the participant’s body, and the outside region covered the other two rows away from the body. To quantify within-subjects and between-subjects variance of proprioception maps, we compute the Pearson correlation coefficients of the error vectors across sessions and between individuals, respectively. The same correlation analysis was also applied to the Euclid distance.

As proprioception errors improved across sessions, we calculated the error reduction as the percentage difference between the first session and the other sessions by 100%× (error_1_ − error_i_)/error_1_, where error_1_ refers to the proprioception error of the first session and error_i_ refers to that of compared sessions (i = 2, 3, 4).

For the trajectory production task, the motor error was defined by the root mean square error (RMSE) between the target trajectory and the participant’s movement trajectory. Each movement trajectory was evenly divided into 30 segments along the y-axis between the start position and the upper edge. The horizontal deviation in the x direction at the cut points of adjacent segments was used to compute the RMSE for each trial:

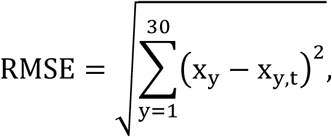

where x_y_ and x_y,t_ is the horizontal ordinate (x value) of the movement trajectory and the target trajectory at the cut points, respectively. For each participant, we computed their average motor error and average proprioceptive error in session 1, and then computed the Pearson correlation between these two baseline performance measures across participants. If the data did not meet the Gaussian assumption, the Spearman’s correlation was computed instead.

Average proprioception errors were compared between sessions or between regions by repeated-measures ANOVAs. Post hoc comparisons between groups were conducted with Bonferroni corrections. The homoscedasticity and normality assumptions were examined before ANOVAs were performed. All dependent variables met these assumptions unless otherwise mentioned. For the data violating homoscedasticity assumptions, Greenhouse-Geisser correction was applied for ANOVAs. For the data violating normal distribution, the natural logarithm function was applied to transform the data into a normal distribution before ANOVAs. One-sample t-tests were used to compare the error reduction percentage of each session with zero. Paired t-tests were used for within-subject comparisons if normality assumptions were satisfied. Otherwise, Wilcoxon t-tests were used. Correlation coefficients were submitted to Fisher’s Z transformation before comparations. All analyses were performed with MATLAB (The MathWorks, Natick, MA) and SPSS version 19 (IBM, Somers, NY). The significance level was set at *p* < 0.05.

### Convolutional neural network classifier

We used the convolutional neural network (CNN) algorithm to investigate to what extent one’s proprioception map was distinguishable from others’. We hypothesize that if the proprioception map is idiosyncratic and stable, a classifier trained by one or two sessions of proprioceptive performance will be able to identify the individual from other individuals based on her/his later performance. Since the data structure of the proprioception map is a matrix similar to a digital image, our CNN classifier was constructed as a typical image classifier (Machine Learning Toolbox, MATLAB 2018b, Natick, MA). The input of the CNN classifier was a 2 × 5 × 20 proprioception error matrix, where the first dimension was the error direction (x and y, two dimensions) at each target position and the other two dimensions representing the coordinate dimensions of the 100 targets. The CNN classifier contained an input layer, a convolution layer, and a normalization layer. A rectified linear unit was applied as an activation function, followed by a drop out layer and a fully connected layer. Finally, a SoftMax function was applied to change the output into the probability of each class. The kernel size of the convolution layer was 3, and the number of output filters was 13. The cross-entropy was used as the loss function, and the Stochastic Gradient Descent with momentum (SGDM) was used to optimize the CNN classifier. The initial learning rate was set at 0.01. The CNN classifier was trained for 250 to 500 epochs according to the size of the training set, and the input sequence was shuffled every epoch.

In Experiment 1, proprioception maps in the first two sessions from each participant served as the training set, and the maps in the third session made up the test set. In Experiment 2, we used session 1, 2, and 3 as the training set and used session 4 as the test set. We also collapsed participants from both experiments to test the classification results: all sessions in Experiment 1 and the first three sessions in Experiment 2 were used. Besides using the first two sessions (session 1 and session 2) to predict the last session (session 3), we also tried to use session 1 to predict session 3, use session 2 to predict session 3, and use session 1 to predict session 2. After training, the CNN classifier was tested by identifying a participant from all participants based on his/her proprioception map in the test set. The performance of the classifier was indexed by the classification accuracy, i.e., the percentage of correctly identified error maps in the test set.

## Results

In Experiment 1, we found that repetitive measurements of proprioception improved subjects’ accuracy of visual matching task. This result is surprising, given that no performance feedback was provided during the measurement. Despite the improvement in the accuracy of proprioception, the spatial characteristics of the proprioception map remained idiosyncratic, as shown by relatively large between-subject variance and relatively small within-subject variance. Further support was that the CNN classifier could identify people with decent accuracy based on her/his proprioception map. Experiment 2 replicated the major findings of Experiment 1 and ruled out the brief performance feedback during the familiarization trials as the cause of improvement in proprioception across sessions.

### Experiment 1

Experiment 1 aimed to examine whether the proprioceptive performance is idiosyncratic and stable across a day. Interestingly, the average proprioception error significantly reduced over the three sessions (*F*(2,50) = 12.368, *p* < 0.001, one-way ANOVA; Fig 2A). The average proprioception errors of three sessions were 3.098 ± 0.776 cm, 2.944 ± 0.767 cm, and 2.420 ± 0.581 cm, respectively (means ± SD, same below). Post hoc pairwise comparisons indicated the proprioception error of the third session was significantly smaller than that of the first session (*p* < 0.001) and the second session (*p* = 0.003). However, there was no significant difference between the first and the second session (*p* = 0.787), which means the proprioceptive accuracy only improved significantly on the second day. The error reductions for the second and third sessions were 3.15 ± 20.95% and 19.05 ± 20.72%, respectively. Only the third session had error reduction that was significantly larger than zero (*t*(25) = 4.657, *p* < 0.001, one-sample t-test). For the trajectory matching task, the average movement error was 2.172 ± 0.595 cm. The movement error did not correlate to the average proprioception error in session 1 (Fig 2B, *r* = 0.267, *p* = 0.187) or session 2 (*r* = 0.295, *p* = 0.143). Thus, the accuracy of proprioception measured by the visual-matching task appears not predictive of the performance of the trajectory production task, though both tasks require accurate localization of the hand in the reachable space with visual targets.

**Figure 2:**
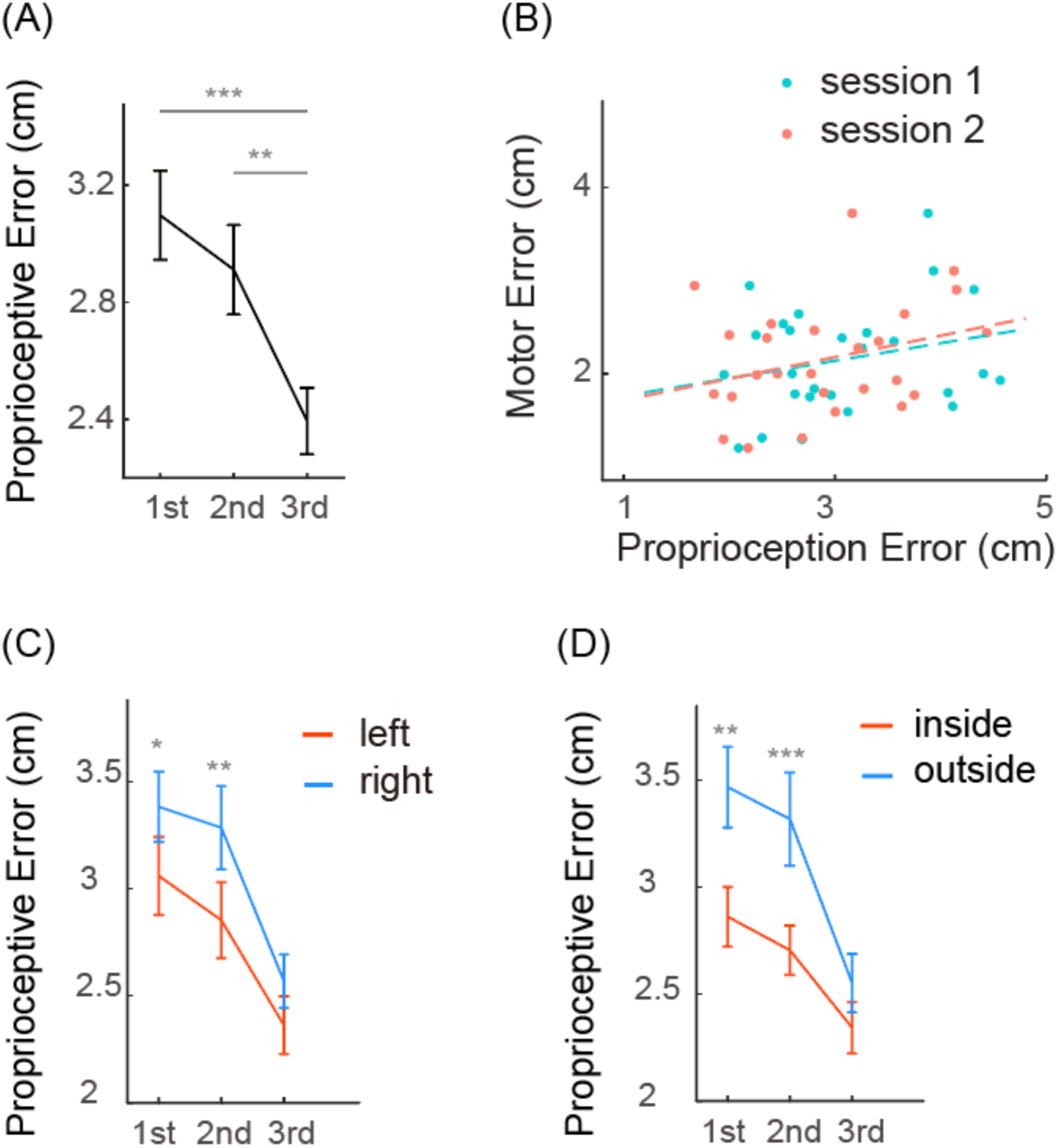
Proprioception error and motor error in Experiment 1. A) Average proprioception error in different measurement sessions. B) Scatter plot of motor errors and proprioception errors from individual participants. The proprioception errors are plotted separately for session 1 and session 2. The dots lines indicate their corresponding linear fits. C) Average proprioception error of the left region and the right region. D) Average proprioception error of the inside region and the outside regions. Error bar denotes SE. \**p* < 0.05; \*\**p* < 0.01; \*\*\**p* < 0.001.

On the group level, the proprioception map showed similar spatial heterogeneity as in previous studies (van Beers et al. 1998; Haggard et al. 2000; Fuentes and Bastian 2009). In the reachable workspace, the proprioceptive error was larger on the right side than on the left side. The proprioception error of the left region were 3.061 ± 0.934 cm, 2.855 ± 0.905 cm, and 2.363 ± 0.690 cm for session 1, 2 and 3, respectively. The proprioception error of the right region were 3.386 ± 0.838 cm, 3.287 ± 0.994 cm, and 2.569 ± 0.637 cm, respectively (Fig 2C. left). The proprioception error of the left region was significantly smaller than that of the right region in session 1 (*t*(25) = -2.587, *p* = 0.016, paired t-test) and 2 (*t*(25) = -2.983, *p* = 0.006, paired t-test), but not in session 3 (*t*(25) = -1.850, *p* = 0.076, paired t-test). On the other hand, the proprioceptive error was larger on the far side of the workspace than on the near side. The proprioception errors of the near region were 2.861 ± 0.771 cm, 2.704 ± 0.588 cm, and 2.341 ± 0.613 cm for the three sessions, respectively. The proprioception errors of the far region were 3.465 ± 0.961 cm, 3.316 ± 1.113 cm, and 2.549 ± 0.696 cm, respectively (Fig 2C., right). Again, the difference between these two regions was significant in session 1 (*t*(25) = -2.992, *p* = 0.006, paired t-test) and 2 (*t*(25) = -5.665, *p*<0.001, paired t-test), but not in session 3 (*t*(25) = -1.835, *p* = 0.078, paired t-test). The improvement from session 1 to session 3 was also larger in the far region (0.916 ± 0.915 cm) than in the close region (0.520 ± 0.767 cm, *t*(25) = -3.506, *p* = 0.002, paired t-test). However, the improvement of the right region (0.745 ± 0.733 cm) and the left region (0.612 ± 0.945 cm) was not significantly different (*t*(25) = - 1.040, *p* = 0.308, paired t-test). In summary, participants performed better in the left region and in the near region when proprioception was measured in the reachable workspace. These regional differences tended to decrease with improvement in proprioceptive errors over successive sessions. It is worth noting that the measurement session did not provide any feedback about their performance. The only occasion that performance feedback was provided was the 16 familiarization trials before session 3.

The error vectors of all participants at 100 target positions were averaged to construct a group-level proprioception error map (Fig 3A). The error map of sessions 1, 2, and 3 shared a certain level of similarity. For example, the error vectors of session 1 generally pointed to the same directions as those of session 2 and 3. For all sessions, most of the error vectors pointed rightwards with larger error magnitudes when more away from the left shoulder.

**Figure 3:**
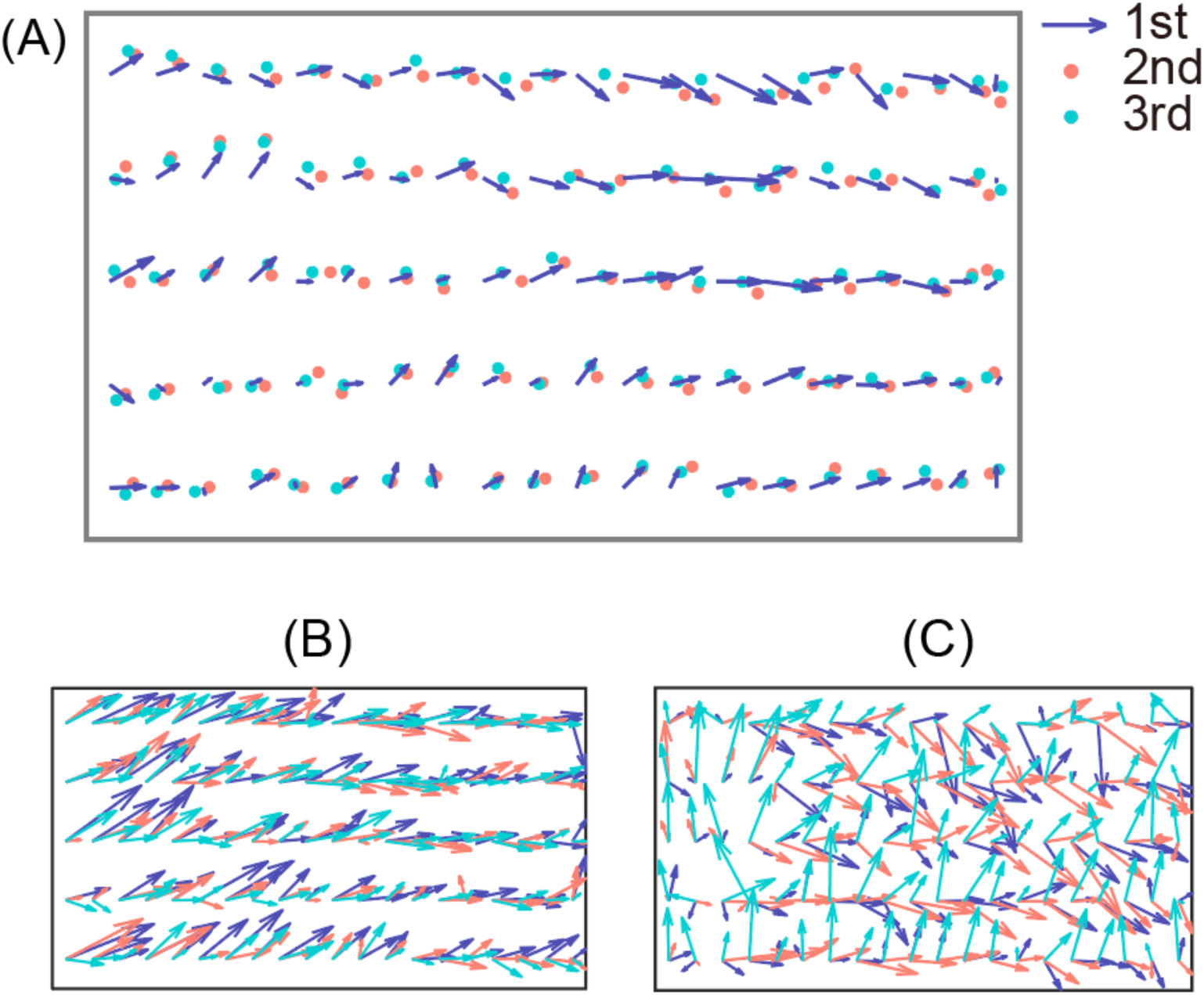
Proprioception maps on the group level and from two selected participants. A) Error map averaged over all participants. The purple arrow denotes the error vector of session 1 with its tail at the target location and its head at the actual hand location. The red and green dots denote the actual hand location in session 2 and 3, respectively. B) Proprioception maps from a typical participant whose error patterns remained similar across measurement sessions. The inter-session correlation coefficient was 0.68, 0.73 and 0.73 for session 1 vs. 2, 2 vs. 3, and 1 vs. 3, respectively. C) Proprioception maps from a typical participant whose error patterns changed dramatically across sessions. The inter-session correlation coefficient was 0.58, -0.04, -0.08, respectively.

To quantitively examine the similarity between proprioception maps, we calculated the correlation of proprioception maps between session 1 and 2, between session 2 and 3, and between session 1 and 3 for each participant. The average correlation coefficients were 0.462 ± 0.216, 0.499 ± 0.196 and 0.412 ± 0.245, respectively. Examining individual participants, we found that 25 (sessions 1 and 2), 24 (sessions 2 and 3), and 23 (sessions 1 and 3) out of the 26 participants showed significant correlations. These results indicate that the proprioception map remained stable across sessions for most participants (see a typical participant in Fig 3B), and only a couple of participants showed large changes across sessions (see a typical participant in Fig 3C). We found that correlation coefficients were significantly larger than zero on the population level (all *t*(25)*s* > 7, *p*s < 10^−7^). To establish a baseline correlation between error maps, we computed all possible pairwise correlations between every two participants (*n* = 26*25 for each of the three session pairs). For example, for the correlation between session 1 and session 2, we calculated the correlation coefficients between the 1st participant’s proprioception map in session 1 with proprioception maps of participants 2 to 26 in session 2, and thus obtained 25 correlation coefficients. The same procedure was applied for each participant, resulting in 25*26 correlation coefficients that characterized the between-subject similarity of proprioception maps. The between-subject correlation coefficients were 0.153 ± 0.251, 0.147 ± 0.224, and 0.139 ± 0.223 for session 1 and 2, session 2 and 3, and session 1 and 3, respectively. These correlation coefficients were also significantly larger than zero due to the large sample size (all *t*(649)s > 15, *p*s < 10^−7^).Importantly, for all three types of pairwise correlations, the within-subject correlation coefficients were significantly larger than the between-subject correlation coefficients (all *t*(649)s > 5, *p*s < 10^−6^, t-test; Fig 4A). Thus, proprioception maps indeed demonstrated cross-session consistency within individuals.

**Figure 4:**
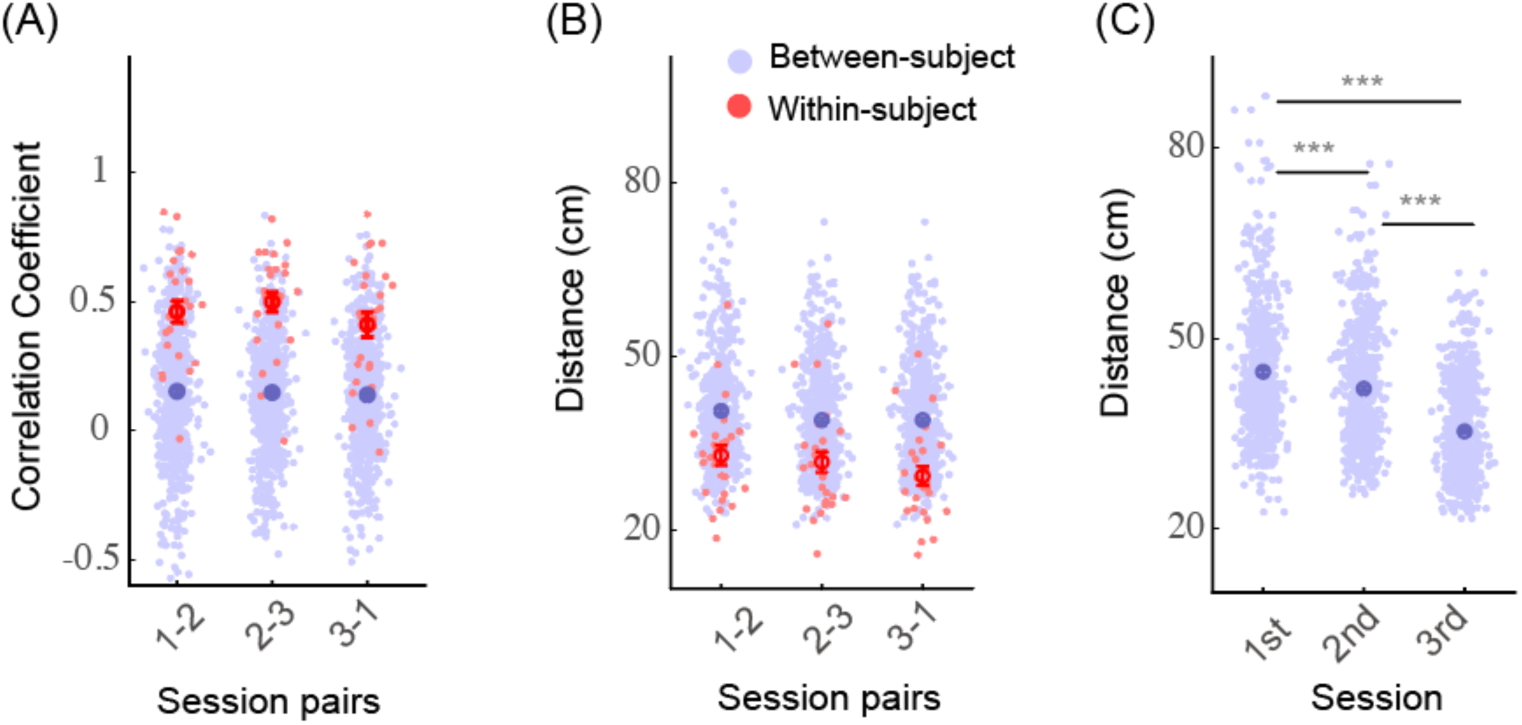
subject-specificity of proprioception error map in Experiment 1. A) Correlation coefficients between session pairs. Blue dotes denote between-subject coefficients. Red dotes denote within-subject coefficients. Error bars denote mean and SE, the same below. B) Comparisons of Euclidean distance between pairs of proprioception maps. C) The Euclidean distance of proprioception maps from each pair of participants within a session. Each blue dot stands for a distance measure between a pair of participants, and the error bar denotes mean and SE. *** *p* < 0.001.

The within-subject and between-subject Euclidean distances between proprioception maps were also compared to evaluate the participant specificity in the same way as the correlation coefficient (Fig 4B). The within-subject distances (mean: 29.4-33.0 cm, SD: 8.2-9.1cm) were significantly smaller than the between-participant distances (mean: 39.0-40.6cm, SD: 8.9-9.1cm) for all three groups (all *Z*s < -4, all *p*s < 0.001, Wilcoxon t-test). In sum, proprioception errors remain idiosyncratic across sessions and days despite the improvement in average proprioception error.

We observed that the between-subject variance declined across time. The distance between the proprioception map of every two participants decreased across three successive sessions (*n* = 650, *Kendall’s W*=0.236, *p* < 0.001, Fig 4C). Post hoc pairwise comparison showed a significant decrease between every two successive sessions (first-second: *Z* = 3.913, *p* < 0.001; second-third: *Z* = 9.391, *p* < 0.001, Wilcoxon t-test), which indicates the idiosyncratic pattern of proprioception might decrease with repetitive measurements.

A convolutional neural network (CNN) classifier was trained and tested with the proprioception maps to perform people identification. The CNN classifier was trained for 350 echoes with the data from the first two sessions and tested with the data from session 3. The training accuracy reached 100%, and the testing accuracy reached up to 73.08% (19/26), which was substantially higher than the chance level (1/26). This means that the classifier was able to correctly identify most individuals by their performance in session 3 on day 2 if their performance on day 1 was provided. From this perspective, the spatial pattern of proprioception error was a person-specific feature even when it changed over time with learning.

### Experiment 2

In Experiment 1, we observed significant improvement of proprioception accuracy across sessions despite that no performance feedback was provided during the measurement. One trivial explanation is that the 16-trial familiarization with feedback before session 3 might serve as a learning session for the visual-matching task. In Experiment 2, we thus canceled the 16-trial familiarization before session 3 to examine this possibility. On day 2, we also added another 16-trial familiarization after session 3 and before session 4 to further examine whether familiarization trials with feedback would lead to the improvement in the proprioception test. Consistent with Experiment 1, the proprioceptive accuracy improved with repetitive measurements (*F*(2.10,42.04) = 4.528, *p* = 0.015, one-way ANOVA; Fig 5A). Post-hoc pairwise comparisons indicated that the proprioception error of both session 3 (*p* = 0.025) and session 4 (*p* = 0.048) was significantly smaller than the first two sessions on day 1. The error reductions of session 2, 3, and 4 were 1.03 ± 18.89%, 10.88 ± 16.43% and 12.69 ± 20.39% respectively (means ± SD; Fig 5B), with the latter two significantly larger than zero (session 2: *t*(20) = 0.258, *p* = 0.799; session 3: *t*(20) = 3.035, *p* = 0.007; session 4: *t*(20) = 2.853, *p* = 0.010, one-sample t-test). The improvement in session 3 confirmed that the improvement observed in Experiment 1 was caused by repetitive measurements as opposed to feedback-based learning in the 16 familiarization trials. Providing familiarization trials with feedback before session 4 did not further improve the performance (*p* = 1.000), further against the possibility of feedback-based learning.

**Figure 5:**
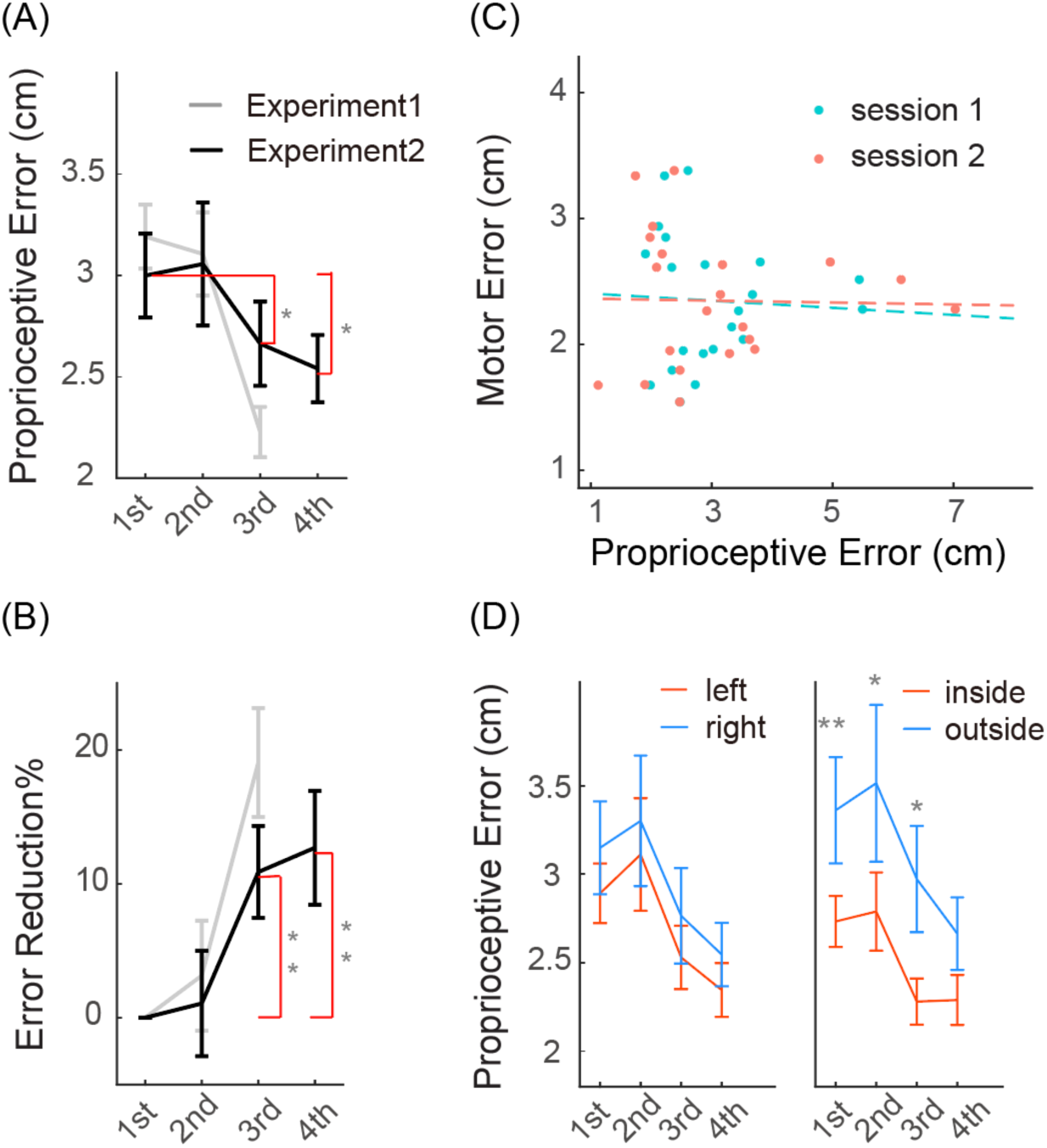
Proprioception error and motor error in Experiment 2. A) Average proprioception error. The black line denotes the average proprioception error for each session. The grey line denotes the corresponding values measured in Experiment 1. B) Reduction of proprioception error reduction as a percentage of error in session 1. The black line and the grey line denote the results from Experiment 1 and 2, respectively. C) Scatter plot of motor errors and proprioception errors from individual participants. The proprioception errors are plotted separately for session 1 and session 2. The dots lines indicate their corresponding linear fits. D) Comparisons of proprioception error between the left and right regions (left), and between the inside and outside regions (right). Error bar denotes SE. \**p* < 0.05, \*\**p* < 0.01.

Experiment 2 also replicated other findings in Experiment 1 (Figure 5). There was no significant correlation between the trajectory-matching error (2.346 ± 0.527 cm, mean ± SD) and the proprioception error in session 1 (*r* = -0.105, *p* = 0.649, Spearman correlation, Fig 5C) or session 2 (*r* = -0.087, *p* = 0.707, Spearman correlation). Comparing average proprioceptive errors in different workspaces, we found that the means of error in the right region was larger than that in the left region in all four sessions, although none of comparisons reached significance (*p*: 0.110 - 0.859, Fig 5D. left, Wilcoxon t-test). The error of the near region was significantly smaller than the error of the far region in the first three sessions (session 1: *Z* = -2.868, *p* = 0.004; session 2: *Z* = -2.103, *p* = 0.035; session 3: *Z* = -2.520, *p* = 0.012, Fig 5D. right, Wilcoxon t-test), but not in session 4 (*Z* = -0.921, *p* = 0.357, Wilcoxon t-test). Similar to Experiment 1, the improvement from session 1 to session 4 was larger in the far region than in the close region (*t*(20) = -2.228, *p* = 0.038), but the improvement was similar between the left region and the right region (*t*(20) = -0.399, *p* = 0.694).

In Experiment 2, we continued to observe that the idiosyncratic pattern of proprioception maps persisted across sessions. For the six session-pairs (session 1 vs 2, session 2 vs 3, session 3 vs 4, session 1 vs 4, session 1 vs 3, session 2 vs 4), the within-subject correlation coefficients had a mean of 0.35-0.548 and a standard deviation of 0.161-0.260. The between-subject correlation coefficients had a mean of 0.070-0.099 and a standard deviation of 0.239-0.291. All the within-subject correlation coefficients were significantly larger than the corresponding between-subject correlation coefficients (all *t*s > 6, *p*s < 10^−5^, t-test, Fig 6A). Furthermore, the within-participant distances (mean: 27.2-36.9 cm, SD: 7.9-12.6 cm) were smaller than the between-participant distances for all six comparison pairs (mean: 40.7-49.9 cm, SD: 13.1-17.3 cm, all *Z*s > 3.3, *p*s ≤ 0.001, Wilcoxon t-test, Fig 6B). Similar to Experiment 1, the between-subject distances within each session decreased over time (*n* = 210, *Kendall’s W* = 0.256, *p* < 0.001, Fig 6C). Post-hoc pairwise comparisons found significant differences between sessions (1st-3rd, 2nd-3rd, 1st-4th, 2nd-4th, all *p*s < 0.001). Thus, the between-subject difference between proprioception maps decreased across days but not within days.

**Figure 6:**
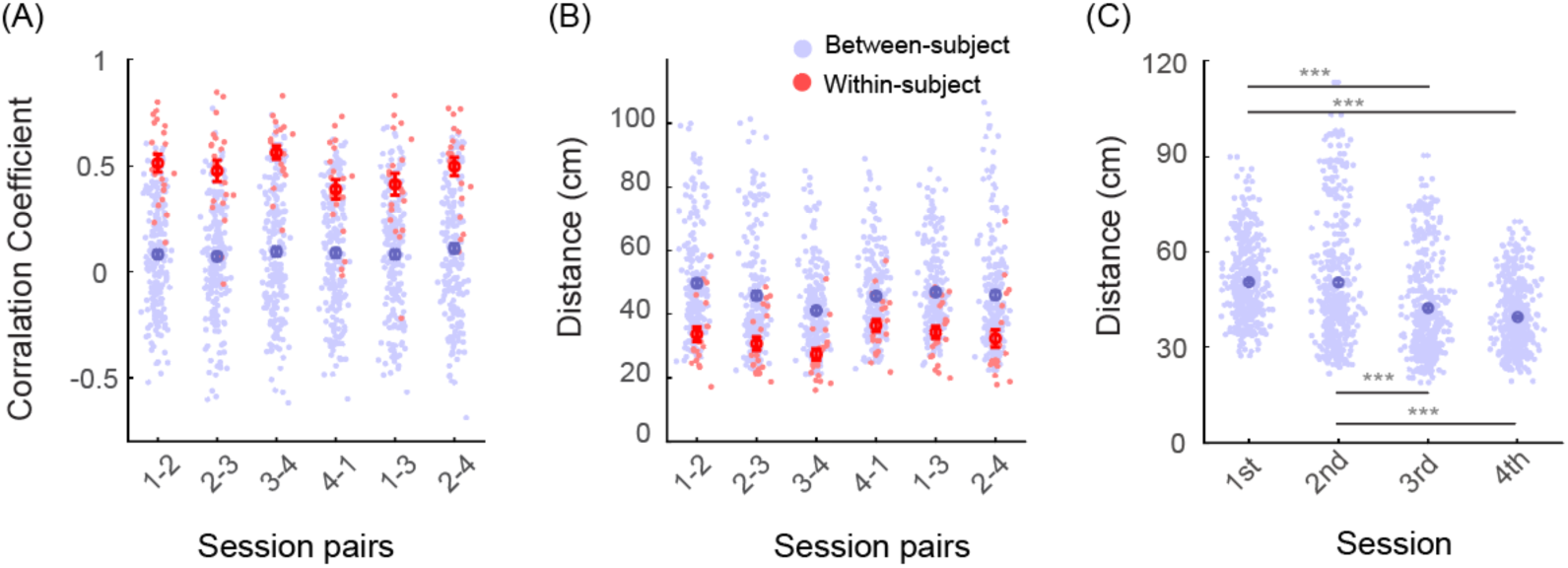
subject-specificity of proprioception error map in Experiment 2. A) Correlation coefficients between session pairs. Blue dotes denote between-subject coefficients. Red dotes denote within-subject coefficients. Error bars denote mean and SE, the same below. B) Comparisons of Euclidean distance between pairs of proprioception maps. C) The Euclidean distance of proprioception maps from each pair of participants within a session. Each blue dot stands for a distance measure between a pair of participants, and the error bar denotes mean and SE. *** *p* < 0.001.

The same CNN classifier, as in Experiment 1, was used to perform people identification based on proprioception maps. To start with, the participants’ proprioception maps from session 1 and 2 made up the training set, and that of session 3 as the test set. After training for 350 echoes, the classifier was able to classify the proprioception from the test set with 76.19% accuracy (16/21). Then, session 1 to 3 were used to train the CNN classifier, and session 4 was used to test it. We obtained a 61.9% testing accuracy (13/21). We also collapsed the data from both experiments to perform people identification with 47 subjects. Using proprioception maps of session 1 and 2 as the training set and third session as the test set, we obtained a testing accuracy of 72.34% (34/47). With this large dataset, we also used data from the first measurement session only as the training set to predict the others. The accuracy could reach 53.19% (25/47) when using session 1 to predict session 2, 55.32% (26/47) when using session 1 to predict session 3, and 61.70% (29/47) when using session 2 to predict session 3. Hence, the CNN classifier could identify individuals with a reasonable accuracy based on a single session of proprioception data. The accuracy can be further improved if an additional session of data was provided as the training data. The overall performance of people identification thus supports that proprioception maps are relatively stable and idiosyncratic among people.

## Discussion

Whether the idiosyncratic pattern of proprioception map persists over time with good within-subject consistency has not been quantitatively investigated in previous research. We used the visual-matching task, a conventional method for measuring proprioception for locating the hand, to repetitively measure proprioception across sessions and across days. We found that 1) humans can improve their proprioception accuracy through repetitive measurements though no performance feedback was given during the measurement, 2) the spatial pattern of proprioception error is subject-specific and remains idiosyncratic across day despite the improvement of accuracy, 3) participants’ proprioception measured in the visual-matching task fails to predict their performance in the trajectory-matching task though both tasks demand accurate location of the hand.

It has been known for long that the error pattern of proprioception varies widely among people (Helms Tillery et al. 1994; Brown et al. 2003a; Smeets et al. 2006; Rincon-Gonzalez et al. 2011), but whether the idiosyncrasy of proprioception maps persists over time has never been tested. We found that the within-subject correlation of proprioception maps between measurement sessions and days was substantially larger than the between-subject correlation. Furthermore, the within-subject dissimilarity between sessions was much smaller than the between-subject one. These findings suggest that the spatial pattern of proprioception map indeed remain consistent over time. Leveraging on the within-subject consistency, a simple CNN classifier could perform people identification based on proprioception maps with fair accuracy. We postulate that subject-specific error pattern might be shaped by individuals’ unique sensorimotor experience in their lifetime since, after all, movement history (Voight et al. 1996; Lee et al. 2003; Forestier and Bonnetblanc 2006) and motor learning experience (Wong et al. 2011, 2012) have considerable influence on one’s proprioception.

The improvement of proprioception without feedback was surprising at first sight. However, although feedback is considered essential for various learning, perceptual learning studies have reported that people can improve without performance feedback in visual perceptual tasks, such as motion-direction discrimination task (Ball and Sekuler 1987) and texture discrimination task (Karni and Sagi 1991). Researchers even have found that the learning rate is similar with and without feedback in a direction discrimination task (Fahle and Edelman 1993). These perceptual improvements are generally attributed to the neural plasticity at the cellular level in the visual system (Petrov et al. 2006). We have similarly found that people can improve their accuracy in the visual-matching tasks with no performance feedback. This finding was observed in two different groups of participants who were tested in two separate experiments. Importantly, our Experiment 2 dropped the 16-trial familiarization trials, thus completely eliminated performance feedback, but continued to observe the improvement of proprioception across days. It is unlikely that this improvement was a result of learning of the task itself since the visual-matching task was easy, and people did not show any improvement between sessions within a day. Hence, we conclude that proprioceptive performance can be improved by repetitive measurements, even when no performance feedback is provided, at least for the widely-used visual-matching paradigm.

For both experiments, the proprioceptive improvement only appeared on the second day, and no improvement was found in session 2 on day 1. Moreover, there was no significant improvement between sessions 3 and 4 on day 2 for Experiment 2. It appears that a night of rest is necessary for the improvement of proprioceptive accuracy. In fact, these findings echo similar findings in other types of perceptual learning where a rest during the night has been shown necessary. For example, in visual studies, one night of sleep is necessary for bringing a performance improvement in a texture discrimination task on the second day (Karni et al. 1994). This improvement is absent if participants are deprived of REM sleep during the night (Walker, Stickgold, Jolesz, & Yoo, 2005). An alternative possibility for our finding is that the manifest of improvement in session 2 might be masked by the trajectory matching task after session 1. Repetitive, active movements could increase the proprioception error in the following measurement session (Kwon et al. 2013). This effect is possibly related to thixotropic behavior of muscles, i.e., intrafusal fibers of muscle spindles become less sensitive to stretch after intensive muscle contraction (Proske et al. 2014). Since muscle spindles play a critical role in proprioception, muscle thixotropy after the motor task could potentially negatively impact the proprioceptive performance measured in session 2. Admittedly, we cannot determine which explanation can account for the lack of improvement within a day, and this issue warrants further investigations.

The visual-matching task used in the present study is a conventional method to measure proprioceptive accuracy (van Beers et al. 1998, 2002; Haggard et al. 2000; Goble et al. 2010; Wilson et al. 2010). If the measurement task itself can reduce the proprioception error, we need to consider its validity as a measurement instrument. For example, a few studies have investigated how visuomotor adaptation of reaching tasks affects proprioception of the hand (Cressman and Henriques 2010; Goble et al. 2010; Ostry et al. 2010; Wong et al. 2011, 2012). These studies typically involve measurements of the proprioception before and after visuomotor adaptation. Our findings suggest that at least part of the changes observed in this kind of study is related to improvement across successive measurements of proprioception. Thus, extra caution is required for the repetitive use of proprioception measurements, such as the visual-matching task.

We found that locating the left hand was more accurate in the left workspace than in the right workspace, and in the area close to the body than away from the body. Furthermore, on the group level, participants perceived their left hand to be more left than its actual position. These spatial patterns of proprioceptive errors were consistent with previous studies (Wilson et al. 2010; Jones et al. 2010). Interestingly, the regional difference of proprioception accuracy tends to diminish over the sessions in both experiments: we observed larger improvement in the far region than in the near region to the body, closing the gap of accuracy between regions. As the overall accuracy improved, the between-subject variance of proprioception maps also decreased. Taken together, we observe a trend that improvement in proprioceptive accuracy reduces the heterogeneity and idiosyncrasy of proprioception maps at the same time. Whether this trend will continue with more learning sessions is worth further investigations.

Our findings indicate that better accuracy in proprioception does not translate to better performance in the trajectory-matching task. The visual-matching task employed here to measure proprioception requires participants to keep their limb stationary with respect to a reference position (Wann and Ibrahim 1992; van Beers et al. 2002; Brown et al. 2003a; Goble et al. 2010). Arguably, this method can only measure participants’ ability to localize their body parts in a static state. The motor performance of our trajectory-matching task, instead, rely on proprioception in a dynamic sense to produce an accurate movement trajectory. The ability to sense the motion of a moving effector is referred to as kinaesthesia (Jones et al. 2010). Indeed, the accuracy of static proprioception and that of kinaesthesia do not correlate well (Grob et al. 2002). Our findings further suggest that an individual’s performance in static proprioception does not predict her/his motor performance that critically depends on accuracy in locating a moving effector.

However, this conclusion appears contradictory to previous findings of the beneficial effect of motor learning on proprioception (Wong et al. 2011) and the beneficial effect of proprioceptive training on motor learning (Wong et al. 2012). We postulate that Wong and colleagues’ findings can be better explained by learning generalization between similar tasks. For example, in their first study, the motor learning task required participants to grasp a handle to steer a cursor towards a visual target (Wong et al. 2011). This task was thus similar to our proprioception measurement task in which participants needed to move to and stay at a visual target with their hand. Their subsequent proprioception measurement was conducted by judging the relative position of a passively located hand, which grasped the same handle, with respect to a visual target in the same workspace. Thus, their motor learning task and proprioception measurement task were similar since both involved locating the hand at the end of a movement relative to a visual target. Similarly, in their latter study, the proprioceptive training was performed by passively moving the hand by the handle to “copy” a target circle (Wong et al. 2012). The subsequent motor learning task was performed by actively copying the same target circle. These two tasks thus involved similar target trajectories and kinesthetic inputs during the movements. It is thus not surprising that both studies found improved performance in one task after learning the other as a result of a possible near transfer of learning between similar tasks. As discussed above, our visual-matching task was different from our trajectory matching task since they relied on different aspects of proprioception and involved different visual targets. We postulate that these differences thus lead to a lack of correlation in performance between the two tasks. The difference between our study and Wong and colleagues’ study also highlights the independence of static proprioception and kinaesthesia.

Our experiments have some methodological limitations that need considerations in future studies. For instance, our visual matching task includes a large number of target positions as a means to cover a large workspace, resulting in a relatively long measurement session (around 20 minutes) and a lack of repetition at each target. Whether these factors affect the precision and accuracy of proprioceptive measurements is unknown. Some of the previous studies chose to two alternative force choices (2AFC) to judge the relative position of their hand to a visual reference position after movement. Arguably, 2AFC gives a better measurement of proprioception though it is more time-consuming for obtaining a proprioception map. We suggest that future study should tradeoff between accuracy and duration of proprioceptive measurements while keeping in mind that proprioceptive measurement itself is a form of perceptual learning.

## Conclusion

Our quantitative approach demonstrates that the spatial pattern of proprioception error is indeed subject-specific and relatively stable across time. The idiosyncrasy of proprioception map can be utilized to identify people with fair accuracy based on individual’s performance in the proprioception measurement task. Notably, we have also found that a conventional proprioception measurement, the visual-matching task, is able to improve people’s proprioception accuracy even when no performance feedback is given. This result suggests that extra caution should be taken in future experiments where repetitive measurements of proprioception are needed. Finally, we have found that proprioceptive accuracy measured with static postures fails to predict the performance of a motor task that requires accurate positioning of a moving hand, suggesting a functional independence between static proprioception and kinaesthesia.

## Acknowledgements

This work was supported by the National Natural Science Foundation of China (31671168, 31622029, 61533001, 31871116).

## Author Contribution

T.W. analyzed data; T.W. prepared figures; T.W. drafted manuscript; T.W. and K.W. edited and revised manuscript; T.W. and K.W. conceived and designed research; T.W. and K.W. interpreted results of experiments; T.W., Z.Z., Y.Y., H.H. and I.K. performed experiments; T.W., Z.Z., Y.Y., H.H., I.K. and K.W. approved final version of manuscript.

